# Lipid modification of DNA nanocages enhances cellular uptake, migration, and *in vivo* uptake

**DOI:** 10.1101/2023.05.06.539685

**Authors:** Krupa Kansara, Ramesh Singh, Pankaj Yadav, Ashutosh Kumar, Dhiraj Bhatia

## Abstract

The extraordinary self-assembling nature of DNA nanostructures and high functionality enables the formulation of DNA nanostructures with multiple chemical and biological molecules. How the whole organisms in native as well as modified; and how stable they are inside take up exactly these modified DNA nanostructures the organisms still remains to be explored. Here we report the fabrication and evaluation of a new conjugate of a cationic lipid, N-[one-(two, 3-dioleyloxy) propyl]-N, N, N-trimethylammonium chloride (DOTMA) and DNA tetrahedron nanostructure (TdN) for the enhanced uptake, stability, bioimaging, and biotherapeutics in cells and zebrafish (Danio rerio) eleuthero embryos as a model organism. We summarise the enhanced uptake potential of TdN-DOTMA conjugate for futuristic biomedical applications such as drug delivery, bioimaging, biosensing, and therapeutics.

## 1. Introduction

DNA nanotechnology is highly regarded due to natural DNA genetic molecules^1^. DNA can potentially be self-assembled to form various designer nanostructures due to their complementary sequence^2^. This new era of DNA nanotechnology is witnessing DNA-based technologies in various research fields including smart and controlled nanodevices, biosensing, bioimaging, and various therapeutics^3^. The unique inheritance feature of DNA makes them a suitable candidate for advanced programmability, stability, and size control^4^. The functional nucleic acids i.e. part of DNA nanostructures can modulate and undergo conformational changes with target molecules and are useful for biosensing and bioimaging^5, 6^. Additionally, in another category of DNA-responsive nanodevices, target molecules can bind to functional nucleic acids and undergo transformations in the presence of a specific environment, for example, DNA nanodevices with specific sequences can form i-motif or triplex structures in an acidic environment for pH sensing^7^. Potassium (K+) and sodium (Na+) ions can be sensed through DNA G-quadruplex^8^. Advantageously, these nanodevices can be restructured and reformed in certain switchable conditions. Various structures such as stems, loops, hairpins, and ligand-linked complexes or cargos can be formed through a single DNA strand by different design approaches^9-11^. The secondary structures of DNA can form via the hybridization chain reaction (HCR) strategy and rolling circle amplification (RCA) strategy^12-14^. DNA nanoengineering allows DNA strands to form three-dimensional (3D) nanostructures through base pairing and programming i.e. DNA tetrahedrons (TdN)^15, 16^. In addition, DNA-walking nanorobots can be formed through self-assembly which carries the payload in pre-decided directions^17, 18^. Furthermore, chemical and biological modifications of TdN nanostructures with RNA, small nucleic acid scaffolds, lipids, aptamers, and RNA inhibitors facilitate the targeted and precise delivery^19-21^. The inherited nature of DNA enables it to construct simple to complex and small to large nanostructures with high specificity, making it a potential platform for various biological applications.

The cellular membrane act as a protective barrier for the vital organelles of cells. DNA nanostructures and cell membranes both possess net negative charges thus the uptake of such nanodevices to cross the cellular membrane is difficult due to electrostatic repulsion. However, few studies have reported the uptake of 2D and 3D nanostructures without any transfection agents^22, 23^. DNA nanostructures uptake is highly dependent on a few factors such as cellular morphology, DNA nanostructures charge, size, and chemical composition. The DNA nanostructures can be conjugated and modified with various molecules possessing regular valences and charges, these phenomena can alter their uptake potential and decide their intracellular fate. Li et al. reported TdN modified with triphenylphosphine (TPP) localized in mitochondria, used for mitochondrial-based imaging, and monitored the aerobic respiration and glycolysis which are mitochondrial-related arbitration^24^. However, the potential question still arises is how exactly these nanostructures are taken up by the whole organisms in native as well as targeted or modified along with how stable they are inside the organisms. To enhance the uptake of such DNA nanostructures for various biomedical applications, rigorous improvement in terms of the formulation of DNA nanostructures are of vital importance.

We reported recently the correlation between various geometries of DNA nanostructures with different cell lines^25^. The cellular uptake study was conducted on the SH-SY5Y, HeLa, SUM-159A, KB3, and RP1 cell lines with TdN, icosahedron (ID), buckyball (BB), and cube DNA nanostructures and results demonstrated that the TdN uptake was significantly high compared to other DNA nanostructures^25^. Earlier studies reveal the more efficient conjugation of lipids and general oligonucleotides enhanced their biomedical applications^26-29^. In the present study, we explored the new horizon by conjugating cationic lipid, N-[1-(2,3-dioleyloxy)propyl]-N,N, N-trimethylammonium chloride (DOTMA) with TdN as lipid plays an essential role as the plasma membrane’s lipid bilayer is providing easy diffusion of TdN-DOTMA via van der Waals interactions. The positively charged DOTMA and negatively charged TdN form a binary system having a positively charged head group and a hydrophobic tail can be a useful strategy to neutralize the negative charge on DNA nanostructures and to enhance the internalization with the lipid bilayer of the cell membrane^26, 27, 29, 30^ (**Figure 1**). Zebrafish has attracted researchers from around the globe due to its transparent embryos, least maintenance, high fecundity, attractive visualization of nanomaterials uptake and growth, and around 70% gene similarities to humans^31, 32^. The intracellular fate and stability of DNA nanostructures such as TdN-DOTMA are not fully understood, the uptake mechanism of these TdN-DOTMA nanostructures in developing zebrafish larvae is of vital importance.

**Figure 1.**
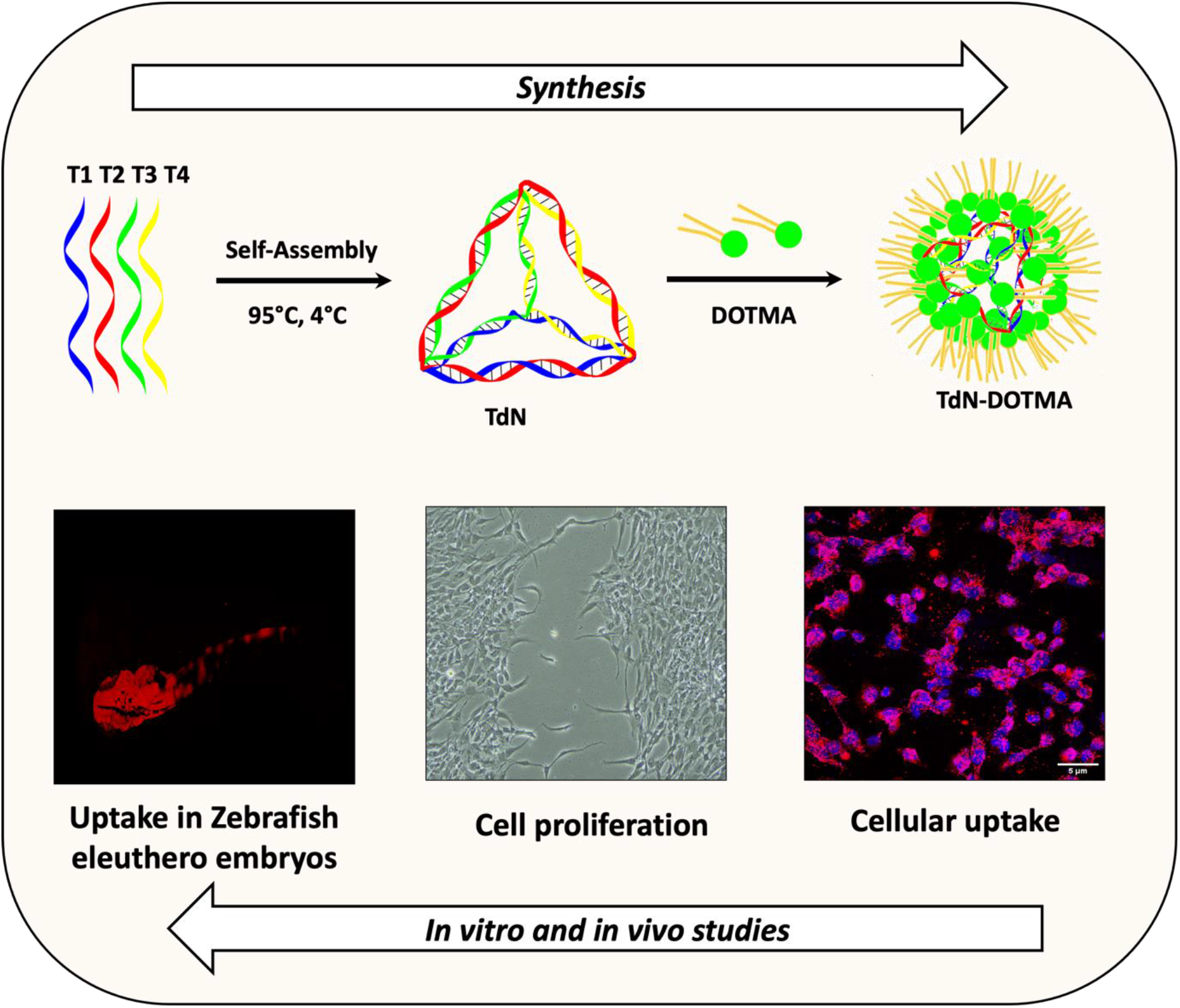
Schematic showing the study design. Cationic lipid DOTMA forms a complex with negatively charged DNA backbone of tetrahedron. This lipid modified DNA nanocages shows enhanced cellular uptake, induces cell migration and enhanced uptake in vivo systems such as zebrafish embryos.

The present study was conducted to investigate the uptake potential in retinal pigment epithelium (RPE1), cell proliferation of TdN-DOTMA nanostructures in a mesenchymal triple-negative breast cancer cell line (SUM-159-A), and zebrafish eleuthero embryos and assess the effect of TdN-DOTMA on the development of zebrafish eleuthero embryos. The eleuthero embryos serve as the ideal *in vivo* model for biological applications^33, 34^. The ratio of TdN: DOTMA (1:100, where DOTMA at concentrations 100 eq.,) was used and optimized for *in vivo* uptake studies based on our recent data^35^. The synergetic effect of TdN-DOTMA on the developmental genes of zebrafish was also analyzed by real-time PCR. The extensive characterization and stability of TdN-DOTMA, and their effect on developing zebrafish eleuthero embryos have provided insights into the biocompatible features of TdN-DOTMA. The cell proliferation potential of TdN-DOTMA has been explored in detail as well.

## 2. Results and Discussion

### 2.1. Characterization of TdN and TdN-DOTMA

The TdN was synthesized using a previously reported protocol and confirmed the formation of TdN by electrophoretic mobility shift assay and Dynamic light scattering (DLS) analysis^36^. The formation of TdN results in retardation in the electrophoretic migration as compared to other monomer nucleotides and smaller assemblies. The average hydrodynamic diameter of the TdN is 12.23 ± 0.25 nm is almost similar to expected. Atomic force microscopy imaging further revealed the formation of homogeneously distributed TdN. Further, the complexation of these TdN with cationic lipids has been carried out by incubation of TdN with 100 equivalent of DOTMA. The co-assembly of the TdN and DOTMA was confirmed by AFM imaging. AFM image of the TdN-DOTMA complex reveals the formation of sphere-like co-assembled structures which are exactly similar to those found in the previously reported^35^ (**Figure 2**).

**Figure 2.**
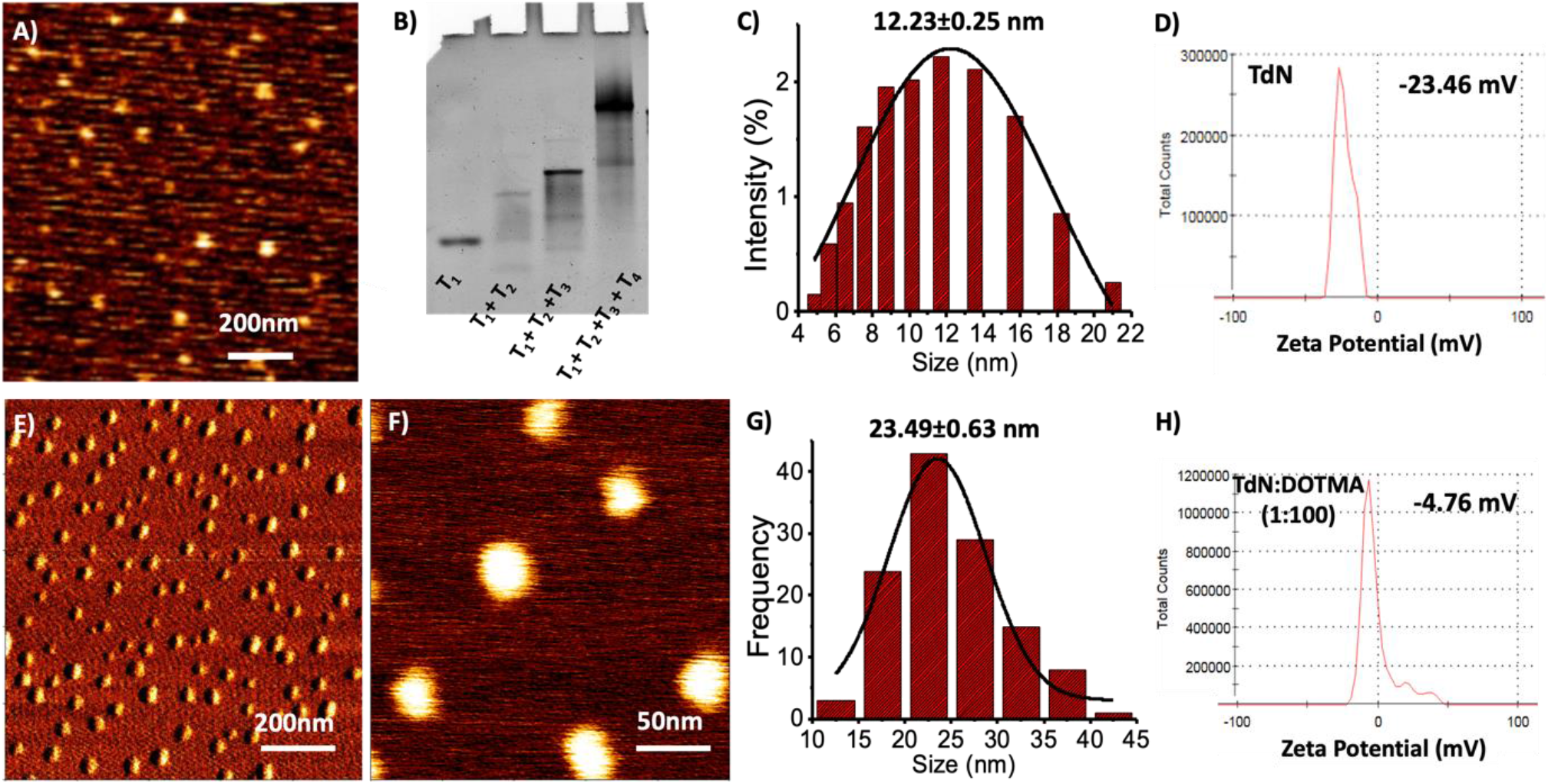
Characterisation of TdN and TdN-DOTMA nanostructures. (A) AFM image of the TdN (scale bar 200nm). (B) Gel electrophoresis mobility shift-based characterization; the delay in mobility indicates the formation of the tetrahedron (T1+T2+T3+T4). (C) DLS results show the hydrodynamic diameter (12.23±0.25 nm) of TdN. (D) zeta potential of TdN (−23.46mV) (E) AFM image of TdN-DOTMA conjugates (scale bar 200nm) (F) A corresponding high-resolution image of TdN-DOTMA assembly (scale bar 200nm). (G) Size distribution bar graph of AFM image E. (G) zeta potential of Td-DOTMA (1:100) complex (−4.76mV). The average size of TD is ∼12.23 nm, and TD-DOTMA is 23.49 nm.

### 2.2. Cellular uptake and proliferation

As in our previous study, we reported that conjugating the cationic lipid with TdN improved its cellular uptake. We performed the cellular uptake experiment for the TdN-DOTMA complex (1:100) and confirmed a significant in cellular internalization. SUM-159-A breast cancer cells were incubated with TdN-DOTMA complex nanostructures for 15 min at 37°C and examined the samples using laser scanning confocal microscopy after staining the nuclei with DAPI. Confocal imaging showed a minimal signal in the red channel for the untreated cells. However, the DOTMA functionalized TdN Cy3 labeled showed a significantly increased fluorescent signal from cells, compared to untreated and non-functionalized TdN (**Figure 3**).

**Figure 3.**
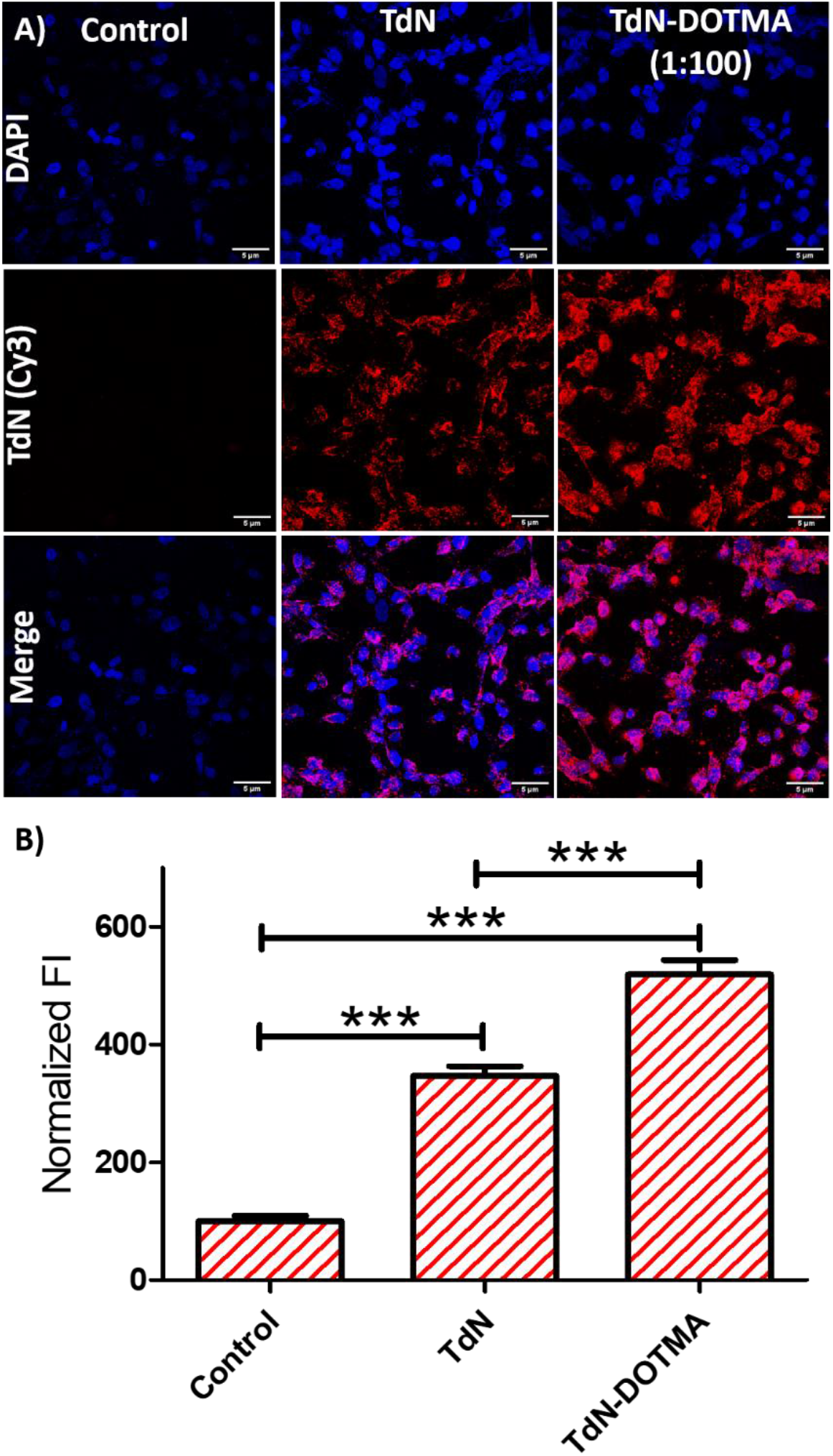
Cellular uptake of TdN– DOTMA (1:100) nanostructures. (A) Confocal imaging of SUM-159-A cells treated with Cy3 labeled TdN and with Cy3 labeled TdN-DOTMA in (1:100) ratios for 30 min. The upper panel (blue channel) represents nuclei stained with DAPI, the middle panel (red channel) represents TdN uptake, and the lower panel is of merged images of the upper two panels. The scale bar is five μm. (B) Quantifying TdN uptake in SUM-159-A cells from panel (A) shows significantly higher uptake of DOTMA modified DNA TdN in cells as compared to control and unmodified TdN. ***P < 0.0001.

The uptake of ligands in cells by specific endocytic pathways is directly connected to cellular processes such as migration and proliferation. The scratch assay was done on RPE1 cells to understand the wound healing capability of TdN: DOTMA (1: 0, 1: 10, 1: 100, and 1: 250). The migration of the cells was noted at time points 0 h, 6 h, 12 h, and 24 h. The migration of the cells was observed in all the conditions of control and TdN: DOTMA (1: 0, 1: 10, 1: 100, 1: 250) at each time point as shown in **Figure 4A**.The migration was studied in control and all the ratios of TdN: DOTMA. At each time point when compared within the control and TdN: DOTMA (1: 0, 1: 10, 1: 100, 1: 250) there is a significant decrease (p< 0.001) in the wound length. A total of 100 wound length points are plotted at each time point in all the conditions of treatment as shown in Figure 4B. The relative average length was calculated (as shown in **Table 1**) and plotted to observe the wound healing. The cell migration was significant in all the treatment conditions: wound length in TdN was 36.64%, in TdN: DOTMA (1:10) was 42.05%, in TdN: DOTMA (1:100) was 50.4%, in TdN: DOTMA (1:250) was 29.07% as compared to RPE1 cells (control) which were 58.08% of the wound length post 24 hours.

**Table 1:**
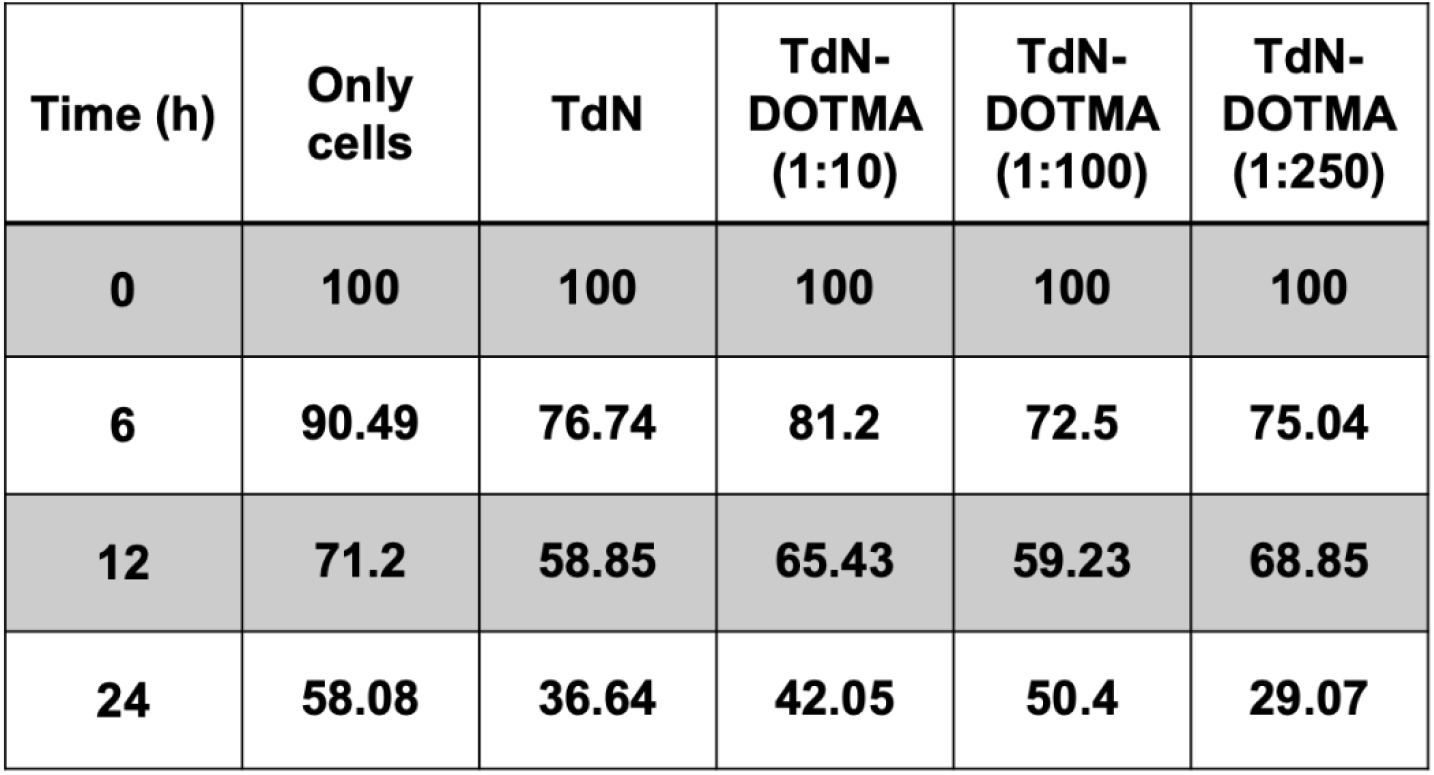
The relative average length (in percentage) of wounds in all the treatment conditions at different time points.

| Time (h) | Only cells | TdN | TdN-DOTMA (1:10) | TdN-DOTMA (1:100) | TdN-DOTMA (1:250) |
| --- | --- | --- | --- | --- | --- |
| 0 | 100 | 100 | 100 | 100 | 100 |
| 6 | 90.49 | 76.74 | 81.2 | 72.5 | 75.04 |
| 12 | 71.2 | 58.85 | 65.43 | 59.23 | 68.85 |
| 24 | 58.08 | 36.64 | 42.05 | 50.4 | 29.07 |

**Figure 4.**
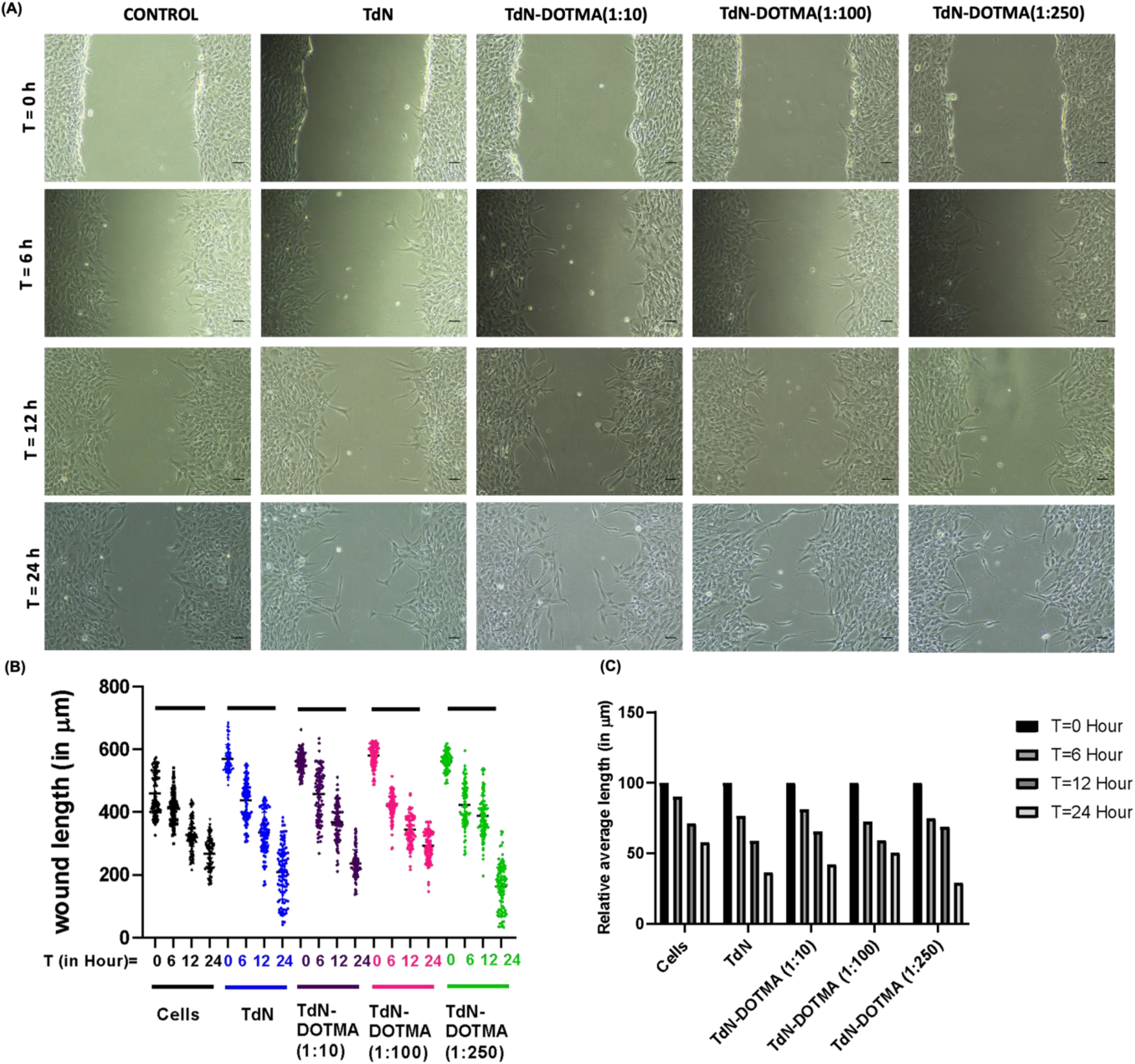
(A) RPE1 cell migration assay in the presence of TdN: DOTMA in the ratio of 1:0, 1:10, 1:100,1:250 as compared to control (RPE1 cells) at time points T = 0, 6, 12, 24 h, the scale bar is 50 micrometer (B) The wound length plot at time points T=0, 6, 12, 24 h in control and TdN: DOTMA in the ratio of 1:0, 1:10, 1:100,1:250. Five images were quantified with 20 length points (in total 100 lengths) in control and TdN: DOTMA in the ratio of 1:0, 1:10, 1:100, and 1:250 at time points T=0, 6, 12, and 24 h. There is a significant difference (p < 0.001) in wound closure in all five conditions (in control and TdN: DOTMA in the ratio of 1:0, 1:10, 1:100,1:250) when compared between time T=0 h and T= 6 h, T=0 h and 12 h, T=0 h and 24 h. (C) The relative average length plot at time points T=0, 6, 12, 24 h in control and TdN: DOTMA in the ratio of 1:0, 1:10, 1:100,1:250 at time points T=0, 6, 12, 24 h. The wound length is 60% in control, 40% in TdN, 48% in TdN: DOTMA (1:10) 50% in TdN: DOTMA (1:100), and 30% in TdN: DOTMA (1:250) when compared at T= 0 h and T=24 h respectively.

### 2.3. Zebrafish eleuthero embryos’ survival rate and heartbeats

To understand the effect of TdN and TdN-DOTMA on survival and heartbeats, zebrafish eleuthero embryos (72 h post-hatching or 144 h after post-fertilization) were exposed to TdN (300 nM) and TdN-DOTMA (1:100). No significant changes were observed regarding survival rate and heartbeats. The reason could be the biocompatible nature of DNA nanostructures. A similar observation regarding the biocompatible nature of TdN has been reported earlier by our group^36^. Our earlier study demonstrated that various concentrations of TdN with different time points on developing zebrafish embryos did not cause any alterations in their survival rate and hatching success^36^.

In this study, we observed that both TdN and TdN-DOTMA exhibited biocompatible nature toward eleuthero embryos post 300 minutes of treatments (**Figure 5a**). The survival rate for TdN and TdN-DOTMA from zero to 300 minutes was the same as for controls. However, alone DOTMA caused a 50% of reduction in survival from 210 to 300 minutes post-treatment. This could be possible due to the chemical nature of DOTMA as cationic lipids such as DOTMA, the cytotoxic effects are mainly determined by the structure of its hydrophilic groups^37^. The heart rate per minute of eleuthero embryos was 150, 142, 126, and 138 for control, TdN, DOTMA, and TdN-DOTMA. There was no significant change in the heartbeat per minute up to 5 hours (**Figure 5b**). The results depict that the TdN and TdN-DOTMA exerted no toxic effect on the survival of the zebrafish eleuthero embryos as compared to the control.

**Figure 5.**
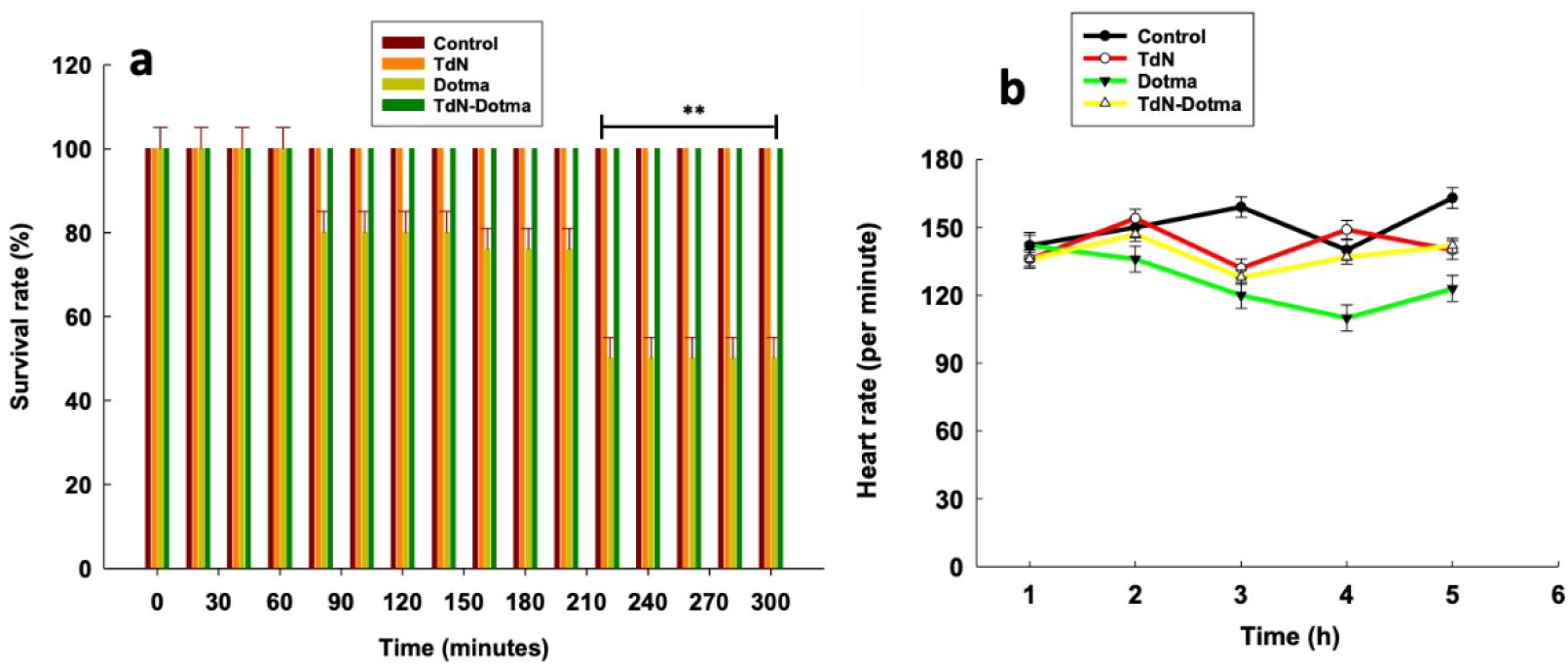
Effects of TdN (300nM) and TdN-DOTMA (1:100) on the survival rate and heartbeats of eleuthero embryos for 5 hours post treatments. (a) Survival rate (b) Heartbeats. Data represent mean ± SEM of three independent experiments. **P < 0.01, when compared with control.

### 2.4. Uptake potential of TdN-DOTMA in Zebrafish eleuthero embryos

Any nanocarrier’s uptake potential is vital for specific biomedical applications. The nanomaterials require some transfecting agents for proper delivery at the cellular level through the plasma membrane however our earlier study has demonstrated that the DNA nanocages were uptaken via endocytic pathways without any transfection agents^36^. With the successful formulation of TdN with DOTMA, we further proceed to analyse and quantify the uptake potential of TdN-DOTMA in zebrafish eleuthero embryos. The eleuthero embryos were exposed to TdN (300nM) and TdN-DOTMA (1:100) for 4 hours of treatment (Figure 3). One of the four oligonucleotides (T4) was labeled with cyanine-5 (cy5) dye at their 5′ ends for tracking purposes. As in our previous report, we demonstrated the significant uptake potential of DNA TdN compared to other geometries of DNA nanocages i.e. buckyball, cube, and icosahedron at several time points such as 4, 6, and 12h post treatments in 72 hours post fertilized zebrafish larva^36^. Therefore, we functionalized TdN with cationic lipid DOTMA to enhance the uptake potential in this study. The uptake of TdN and TdN-DOTMA was quantified using a laser-scanning confocal microscope. The signal for TdN-DOTMA was significantly higher compared to alone TdN in eleuthero embryos post 4 h treatment. Moreover, the physiological process of intracellular transport and the particular qualities of the subcellular structures determine the accuracy of targeted organelles *in vivo*. Precise targeting at the organelle level using fluorescently labeled DNA nanodevices has showed the utility of DNA nanostructures for imaging organelles for diagnostic applications; the same can be investigated for therapeutic applications. The development of DNA nanodevices as diagnostic and therapeutic tools may be aided by understanding the laws of proper transmission between organelles, cells, and tissues. Additionally, our uptake studies indicated that lipid-modified TdN enhanced the uptake and internalization in zebrafish eleuthero embryos. The confocal image of eleuthero embryos clearly showed the uptake and internalization of the TdN-DOTMA nanostructures. Quantification analysis of cy5 signal intensity of TdN-DOTMA exerted the significant uptake with 100 eq. DOTMA compared to alone TdN (**Figure 6**).

**Figure 6.**
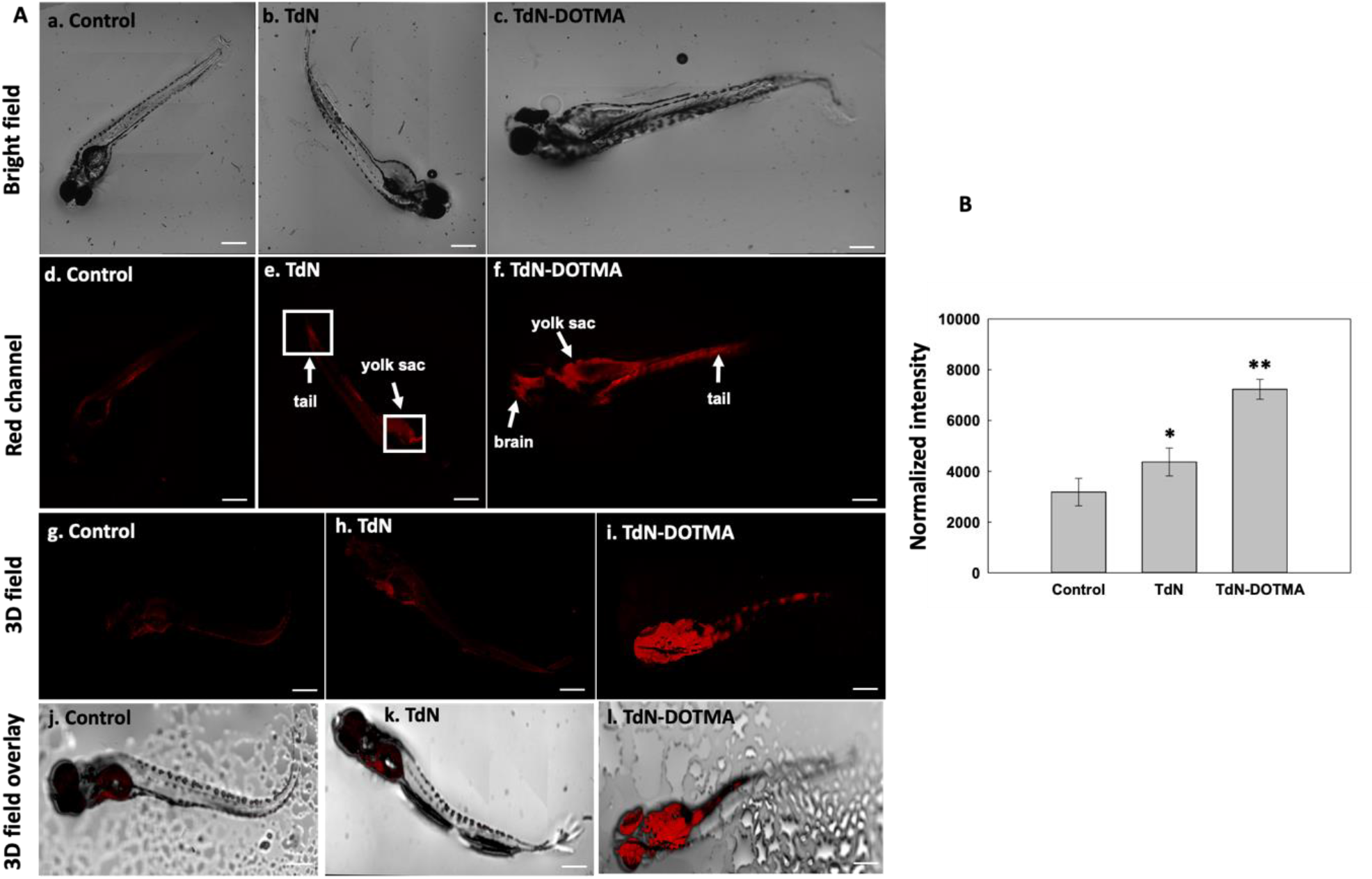
Uptake analysis of TdN and TdN-DOTMA in zebrafish eleuthero embryos. (A) Uptake of TdN and TdN-DOTMA in zebrafish eleuthero embryos post 4 h treatments. (a-c) Brightfield images of zebrafish eleuthero embryos post 4h treatment: (d–f) Red channel images of zebrafish eleuthero embryos post 4h treatment: (g-i) 3D field images of zebrafish eleuthero embryos post 4h treatment: (j-l) 3D field overlay images of zebrafish eleuthero embryos post 4h treatment (B) Fluorescence intensity and quantification analysis of the uptake of TdN and TdN-DOTMA in zebrafish eleuthero embryos post 4 h treatments. The scale bar is set at 50 μm. **Statistically significant p-value. (p < 0.01). *Statistically significant p-value (p < 0.05). 10 eleuthero embryos per condition were quantified.

### 2.5. Effect of TdN-DOTMA on gene expression

Several genes involved in the development of zebrafish eleuthero embryos were targeted to link the developmental changes since the eleuthero embryos exposed to TdN and TdN-DOTMA have a biocompatible effect on survival rate and heartbeats. 12 distinct genes involved in cardiovascular development, dorsal and ventral axis construction and brain network development were targeted. The eleuthero embryos exposed by two groups for 4 h post treatments were studied. For each group, two distinct controls devoid of DNA nanostructures were used. The housekeeping gene was GAPDH (Figure 7).

**Figure 7.**
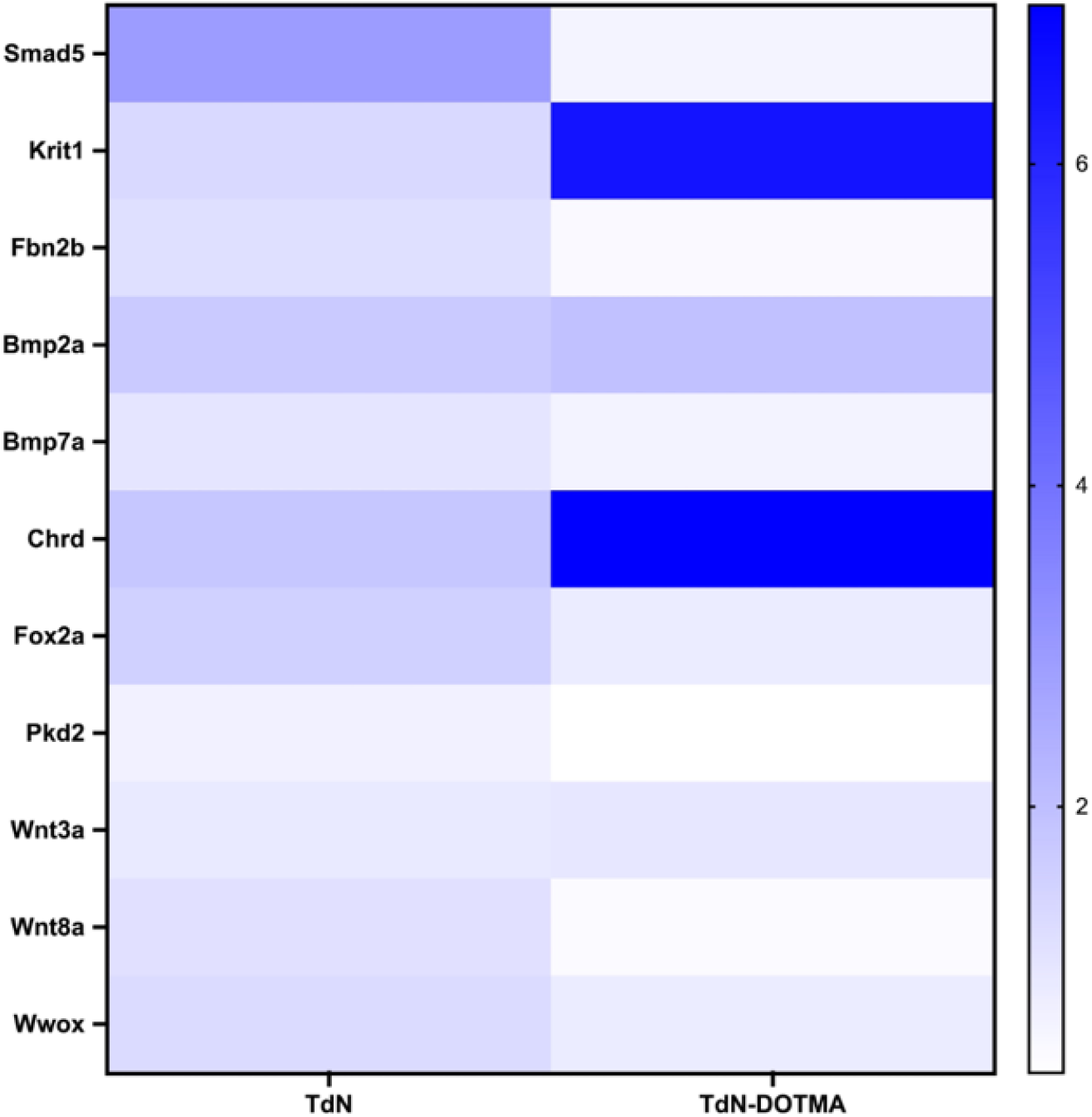
Heat map depicting fold change in relative gene expression of zebrafish eleuthero embryos exposed to TdN (300nM) and TdN-DOTMA (1:100) for 4 h post treatments.

The smad5, chrd, and foxa2 genes, which are primarily in charge of the early development of zebrafish larvae, were evaluated and quantified in this study. According to Higashijima et al., the foxa2 gene is linked to the development of the zebrafish floorplate^38^, and Kansara et al. reported that the chrd gene plays a crucial role in the dorsal-ventral patterning of larvae with the help of TGF-β protein^32^. Smad5 gene expression was increased in TdN-exposed embryos; however, TdN-DOTMA-exposed embryos exhibited normal gene expression compared to the control, which showed the more biocompatible nature of TdN-DOTMA. The results demonstrated the normal expression profile of the smad5 gene except in TdN directly co-related the proper dorsoventrally patterning, which was later observed as healthy eleuthero embryos in TdN-DOTMA treated group post 4 h of treatment.

The bone morphogenetic protein (BMP) signalling pathway is another connection between the smad5 gene and this pathway^39-42^. One of the crucial BMP pathway genes, Bmp7a is in charge of zebrafish embryo and larval proper dorsoventrally development, which results in average growth. To activate SMADs, it typically binds to type one and type two BMP receptors on the cell surface^31, 38, 43-45^. The results demonstrated that TdN-DOTMA exposed embryos’ gene expression did not change significantly compared to the control group for Bmp2a and Bmp7a genes. The expressions were 2 and 2.2 folds for TdN and TdN-DOTMA respectively for Bmp2a genes and 1 and 0.69 folds for Bmp7a (**Figure 7**).

This finding is supported by the survival rate statistics and heartbeat measurements. As the exposure duration for TdN and TdN-DOTMA increased, neither the eleuthero embryos’ deformities nor mortality increased. This might be a result of DNA nanostructures’ biocompatibility, which could be shown in our earlier studies’ survival rates and morphological alterations^36^.

The szl gene and TGF-protein are linked to the chordin (chrd) gene, which is in charge of dorsoventrally patterning ^32, 40^. The dorsalizing activity of the szl gene is related to the reduced production of chrd, which functions as a BMP antagonist. TdN-treated eleuthero embryos exhibited 2-fold increase of chrd gene post 4 h of treatment however, the over-expression of chrd gene was observed in the TdN-DOTMA-treated eleuthero embryos post 4 h treatment. The overexpression of chrd gene might inhibit the BMP pathway as BMP/Chordin has shown antagonist activity. However, our survival rate and malformation observations (data not shown) did not lead to any defects in growing eleuthero embryos.

The Foxa2 gene functions as a transcription factor for the trim46a, which is in charge of the midbrain and MHB (midbrain-hindbrain boundary) and is connected to the development of the floor plate^18, 32, 38^. Developmental problems in the midbrain and MHB as well as aberrant floor plate formation are linked to decreased or increased expression of the foxa2 gene^31^. In all the treatment groups (TdN and TdN-DOTMA) for 4 h treatment, the foxa2 gene expressions were 1.8 folds and 1 respectively. This trend again exhibited the more biocompatible nature of TdN-DOTMA (**Figure 7**).

The Fbn2b gene, also known as the fibrillin-2b gene, plays a vital role in heart morphology and notochord morphogenesis and Krit1 stabilizes the cardiovascular junctions by interacting with other proteins; this gene’s reduced expression or upregulation can lead to severe cardiovascular defects in the zebrafish larva^31, 46^. The expressions of the fbn2b gene in TdN and TdN-DOTMA exposed eleuthero embryos were 1 and 0.5 folds respectively compared to the controls. However, the expressions of the Krit1 gene in TdN and TdN-DOTMA exposed eleuthero embryos were 1 and 6.4 folds compared to the controls. The increased trend of expression of the Krit1 gene in TdN-DOTMA-exposed eleuthero embryos should show some abnormalities in eleuthero embryos however, this observation did not support the heart rate results discussed previously.

Wwox gene is a tumor suppressor gene and is mainly involved in organ development. The under or over-expression profile of the wwox gene can lead to paracardial edema, leading to larval death within a week ^47^. The expression profile of the wwox gene for TdN and TdN-DOTMA exposed eleuthero embryos was 1 and 0.8 folds compared to controls. This result is clear evidence that TdN and TdN-DOTMA did not affect healthy eleuthero embryos and exhibited the biocompatibility nature of DNA nanostructures. The Pkd2 gene is involved in the tail formation, curvature, and mesoderm patterning of zebrafish embryos in their early development ^48, 49^. The malformation data (data not shown) did not exhibit any curly tail or improper development of tail in eleuthero embryos exposed to TdN and TdN-DOTMA and the expression profile of Pkd2 supported the claim as the gene expression was 0.8 and 0.6 folds compared to controls.

In the early stages of zebrafish development, the Wnt signaling system is crucial for body patterning ^50^. Wnt3a and Wnt8a are expressed in the blastoderm’s edges and, in the tail bud, later in development. Pre-somatic mesoderm cells prematurely differentiate into somites when wnt activity is inhibited. As a result, tailbud deformity results from the absence of undifferentiated cells and is caused by the termination of the expression of a specific gene, wnt8a, which is heavily expressed in the tailbud ^51^. Until the 4 h of treatments of TdN and TdN-DOTMA, both wnt3a and wnt8a were not downregulated significantly. The expressions of wnt3a and wnt8a were 1, 1.1, 1, and 0.5 folds compared to controls for TdN and TdN-DOTMA respectively.

## 3. Discussions and Conclusions

We demonstrate here that lipid modification of DNA nanodevices such as TdN-DOTMA conjugates have much more potential for biological uptake than TdN alone. It was also observed that the negative charge of a phosphate ion on the DNA backbone and the positive charge of the ammonium cationic head of the cationic lipid DOTMA were electrostatically attracted to one another and made stable conjugates which have more cellular internalization in SUM-159-A cells than TdN. Similarly, TdN-DOTMA (1:100) exhibited significant migration in non-cancerous RPE1 cells compared to bare TdN. This was equally reflected in terms of their in vivo uptake in zebrafish embryos. Gene expression studies in zebrafish upon treatment with DOTMA-TDN complexes suggest that expressions of most of the developmental genes such as cardiovascular development, dorsoventral axis development, tail formation, and floorplate development exposed to TdN and TdN-DOTMA for 4h post treatments did not decrease and induced any abnormalities in developing zebrafish eleuthero embryos. Here, we described an effective technique to improve the cellular and *in vivo* uptake of DNA nanostructures by functionalizing with a cationic lipid, and that it can be applied to various nano carrier systems for effective internalization in various biological systems for multiple targeted biological and biomedical applications like targeted biosensing, delivery, therapeutics, etc.

## 4. Materials and Methods

### 4.1. Synthesis of TdN and TdN–DOTMA complexation

Previously reported one-pot synthesis was sued for the synthesis of DNA tetrahedrons nanostructures. Four complimentary single-stranded DNA oligos (T1, T2, T3, and T4) in equimolar concentration and 2mM MgCl_2_ were mixed and performed a thermal annealing reaction in a PCR (polymerase chain reaction) instrument with cycling conditions from 95°C to 4°C temperature. The reaction mixture was first heated at 95°C for 10 min and then gradually cooled to 4°C. Cyanine-3 (cy3) and cyanine-5 (cy5) labeled oligonucleotides were used for imaging purposes. A mixture of DNA TdN and DOTMA in a (1:100) molar ratio was incubated for 3h for the functionalization of TdN-DOTMA complexation.

### 4.2. Characterization of TdN and TdN-DOTMA complexes

#### 4.2.1. Electrophoretic mobility shift assay (EMSA)

An electrophoretic mobility shift assay (EMSA) was performed using NativePAGE. 10% polyacrylamide gel was prepared to study the tetrahedral structure formation. The gel was run at 90 V for 90 min then the gel was stained with an EtBr stain and visualized using the gel documentation system (Biorad ChemiDoc MP Imaging System).

#### 4.2.2. Dynamic light scattering (DLS) and zeta potential

The dynamic light scattering (DLS) for hydrodynamic size and zeta potential measurements for charge distribution on TdN were performed with a Malvern analytical Zetasizer Nano ZS instrument. The data were then plotted on OriginPro software followed by a Gaussian fit.

#### 4.2.3. Atomic force microscopy (AFM)

The morphometric characterization of the TdN and TdN-DOTMA complex was performed by atomic force microscopy (AFM). The samples were prepared by following established lab protocols. 5-10 μL aliquots of TdN and TdN-DOTMA (1:100) were spread over a freshly cleaved mica surface and dried. The imaging was performed with Bruker NanoWizard Sense + Bio AFM installed at IITGN Gandhinagar, Gujarat. Particle size distributions in AFM images were measured using ImageJ imaging software and the data acquired were plotted using OriginPro software.

### 4.3 Cellular internalization and proliferation

For the cellular uptake experiment, SUM-159-A cells were used. SUM-159-A cells were maintained in HAMS-F12 complete media containing 10% fetal bovine serum and antibiotic at 37°C and 5% CO_2_ in a humidified incubator. Approximately 10^5^ per well cell counts were seeded on a glass coverslip in a 6-well plate overnight. The seeded cells were washed with 1X PBS buffer thrice and then incubated in serum-free media for 30 min at 37°C and 5% CO_2_ in a humidified incubator before treatment. They were treated with TdN nanostructures and TdN-DOTMA complex (1:100) nanostructures. The treated cells were fixed for 15 min at 37°C with 4% paraformaldehyde and rinsed thrice with 1×PBS.

The wound healing property of TdN: DOTMA (1:0, 1:10, 1:100, 1:250) was studied on RPE1 cells. The cells were cultured and maintained in DMEM complete media in a T25 flask. For the experiment, cells were trypsinized and seeded in a 6-well plate, 24 hours before the experiment. A scratch was made at the start of the experiment using a pipette tip (10 μL). The cells were washed 3 times with 1X PBS and then treated with DMEM serum -free media containing TdN: DOTMA. The cells were checked under a microscope (10x) to visualize their attachment and spreading before the experiment. The cell migration was visualized under the microscope at different time points (t= 0, 6, 12, 24 h).

### 4.4. Zebrafish maintenance

The zebrafish used in this study was of the Assam wild-type strain and was grown in the lab from embryo to adult stage. The husbandry and maintenance of zebrafish were executed according to our previously published protocol ^32^. Briefly, The lab conditions were maintained according to the laboratory conditions explained on ZFIN (Zebrafish Information Network), including a 14 h light / 10 h dark cycle at a temperature of 26 – 28°C. The fish were kept in a 20 L tank with aeration pumps. The water of the tanks was prepared by adding 60 mg/L sea salt (Red Sea Coral Pro salt). The various parameters were maintained to mimic the natural environment i.e., pH (7-7.4), conductivity (250 – 350 μS), TDS (220– 320 mg/L), salinity (210–310 mg/L), and dissolved oxygen (> 6 mg/L) using a multi-parameter instrument (Model PCD 650, Eutech, India). The zebrafish were fed brine shrimp (live artemia) and basic flakes (Aquafin). They were fed the artemia twice and flakes once a day. The breeding setup was prepared in the lab with a ratio of 3 females and 2 males in the breeding chamber. Post breeding, the embryos were collected into E3 medium in a sterile petri dish and kept in a BOD incubator (MIR -154, Panasonic, Japan) at 28.0°C. The embryos were raised in the same medium for three days and then the healthy larvae were used for the experiment.

### 4.5. Survival rate and heat beats analysis

Survival rate studies were conducted by distributing 15 eleuthero embryos per well in a six-well plate (Corning, NY, USA). Volume in each well was made up to 5mL. One well was designated as a control in each group without any nanostructures exposure. The final concentration of TdN was 300 nM and TdN-DOTMA was in a 1:100 ratio (DOTMA at concentrations 100 eq.,).

Our set-up consists of a stereo-zoom microscope utilized to measure the heartbeats of eleuthero embryos manually. Heart rate studies were conducted by distributing 15 eleuthero embryos per well in a six-well plate (Corning, NY, USA). Volume in each well was made up to 5mL. One well was designated as a control in each group without any nanostructures exposure. The final concentration of TdN was 300 nM and TdN-DOTMA was in a 1:100 ratio.

### 4.6. *In vivo* uptake of TdN-DOTMA in Zebrafish eleuthero embryo

*In vivo*, uptake assays were performed according to the Organization for economic cooperation and Development (OECD) guidelines. At 72 h after hatching, corresponding to 144 h post-fertilization (hours post-fertilized), the dead embryo was removed and the remaining embryos were placed in six-well plates (Corning, NY, USA) with 15 eleuthero embryos in each well. Two groups of eleuthero embryos were treated with TdN and TdN-DOTMA at concentrations of 300nM and 1:100 (TdN: DOTMA) ratio and incubated for 4 hours. One well was designated as a control in each group without nanoparticles. Post-treatment, the medium was replaced with fresh E3 media, and eleuthero embryos were washed to remove the excess TdN and TdN-DOTMA and fixed with a fixative solution (4% PFA) for 2 minutes. Post fixation, the eleuthero embryos were mounted with mounting solution Mowiol and allowed to dry for further confocal imaging analysis.

### 4.7. Confocal Imaging and Processing

The confocal imaging of fixed cells (63x oil immersion) and fixed tissues/embryos (10x) was performed using Leica TCS SP8 confocal laser scanning microscope (CLSM, Leica Microsystem, Germany). The pinhole was kept in 1 airy unit during imaging. Image quantification analysis was performed using Fiji ImageJ software. For the quantification analysis, whole cell intensity was measured at maximum intensity projection, the background was subtracted, and measured fluorescence intensity was normalized against unlabelled cells. A total of 40-50 cells were quantified from collected z-stacks for each experimental condition. A total of 10-12 eleuthero embryos from each sample were quantified and 4-5 z-stack were taken from each sample. For comparison of cellular uptake of TdN and TdN-DOTMA, the fluorescence intensity from eleuthero embryos was normalized concerning the blank control where the signal from the nontreated eleuthero embryos was considered as 100.

### 4.8. Total RNA isolation

The total RNA isolation was executed through our earlier protocol^31^. In brief, 20 eleuthero embryos were exposed to 2 groups (TdN and TdN-DOTMA) until 4 h. Total RNA isolation was conducted by RNAeasy mini kit (Qiagen 74104). RNA yield and integrity were determined using the SYNERGY-HT multiwell plate reader (Bio-Tek, USA) using Gen5 software.

### 4.9. Gene expression analysis

First-strand cDNA synthesis was performed with total RNA isolated from 20 eleuthero embryos per replicate with a Maxima First Strand cDNA Synthesis Kit (ThermoFisher Scientific, USA) as per the manufacturer’s protocol. A total of 20 eleuthero embryos were exposed to 2 groups (TdN and TdN-DOTMA) until 4 h then real-time PCR was performed using cDNA from each replicate and gene-specific primers with a powerup™ sybr® green master mix (ThermoFisher Scientific, USA) in the Quantstudio 5 (ThermoFisher Scientific, USA) according to the manufacturer’s protocol. The signal output for each gene was normalized to the level of glyceraldehyde 3-phosphate dehydrogenase (GAPDH) to generate a relative expression ratio. Three independent experiments were performed for each gene, and the fold changes were averaged for a given sample.

### 4.10. Statistical analysis

All the experiments were performed in triplicates, and for the statistical analysis, the results from the experiments were compared with the corresponding control. The results were expressed as the mean ± standard error (SE). The statistical analysis was performed using Sigma plot® 10.0, GraphPad Prism 9.0, and OriginPro software. The image quantification was done using Fiji ImageJ software (NIH).

## Acknowledgements

We sincerely thank all the members of the DB group for critically reading the manuscript and for their valuable feedback. KK thanks SERB, GoI, for National Postdoctoral Fellowship. RS thanks Gujcost-DST and IITGN-MoE, GoI for Postdoctoral fellowship, PY thanks IITGN-MoE, GoI for Directors PhD fellowship. DB thanks SERB, GoI for Ramanujan Fellowship, IITGN for the start-up grant, and DBT-EMR, Gujcost-DST & GSBTM for research grants. Imaging facilities of CIF at IIT Gandhinagar are acknowledged.

